# Overexpressed microRNA-141-3p enhance proliferation via targeting PLAG1 in non-diabetic macrosomia

**DOI:** 10.1101/338335

**Authors:** Dan Guo, Hua Jiang, Yiqiu Chen, Jing Yang, Ziqiang Fu, Jing Li, Xiumei Han, Xian Wu, Yankai Xia, Xinru Wang, Liping Chen, Qiuqin Tang, Wei Wu

**Affiliations:** State Key Laboratory of Reproductive Medicine, Institute of Toxicology, Nanjing Medical University, Nanjing, China; Key Laboratory of Modern Toxicology of Ministry of Education, School of Public Health, Nanjing Medical University, Nanjing, China; Department of Preventive Health Branch, Nanjing Jiangning Hospital Affiliated to Nanjing Medical University, Nanjing, China; Department of Gynecoloy, The Affiliated Obstetrics and Gynecology Hospital of Nanjing Medical University, Nanjing Maternity and Child Health Care Hospital, Nanjing, China; Department of Public Health, Xuzhou Medical University, Xuzhou, Jiangsu, China; Interdisciplinary Toxicology Program, University of Georgia, Athens, Georgia, USA; Department of Gynecology and Obstetrics, Second Affiliated Hospital of Nantong University, Nantong, China; Department of Obstetrics, The Affiliated Obstetrics and Gynecology Hospital of Nanjing Medical University, Nanjing Maternity and Child Health Care Hospital, Nanjing, China; National Institute of Environmental Health Sciences, National Institutes of Health, Department of Health and Human Services, Research Triangle Park, USA.

**Keywords:** macrosomia/microRNAs/miR-141-3p/placenta/PLAG1

## Abstract

Several studies have shown microRNAs (miRNAs) could regulate the placental development, yet the role and mechanism of miRNAs in the development of non-diabetic macrosomia (NDFMS) remains unclear. The key miRNA that abnormal expressed in NDFMS placentas was screened out by miRNA microarray and verified using qRT-PCR in 91 subjects. The effects of the key miRNA were verified by proliferation assay and invasion assay in HTR-8/SVneo cell, and also in pregnant C57BL/6J mice. miR-141-3p was determined as the key miRNA with the most significant difference, which could promote the proliferation and invasion by regulating the expression of target gene *PLAG1*. Overexpression of *PLAG1* could reverse the effect of cell proliferation and invasion ability caused by miR-141-3p overexpression. Significant difference in fetal birth weight was observed between the control group and treated group with miR-141-3p agomir in late pregnancy, but not in early pregnancy. This study revealed miR-141-3p could increase the proliferation of placenta to participate in the occurrence and development of NDFMS through regulating *PLAG1* expression.

## Introduction

As one of the most common perinatal complications of pregnancy, non-diabetic macrosomia (NDFMS) has a serious impact on the long-term health of the baby. To explore the potential causes and mechanisms of NDFMS is significant to reduce the incidence of NDFMS and improve perinatal maternal, child health, and infant long-term health.

Neonatal birth weight is associated with a variety of factors, including genetic factors, gestational nutrition, endocrine and placental function, etc (1, 2). The placenta acts as a bridge between mother and fetus, and can also protect fetus from the maternal immune system attack and secrete pregnancy-related hormones and growth factors (3). Therefore, placental development and abnormal function may have an important impact on the growth and development of the fetus (4, 5). Several studies have shown that microRNAs (miRNAs) participate in all stages of embryonic development and pregnancy. miRNAs are involved in placental tissue development, and play an important role in pregnancy (6, 7). However, the underlying mechanism is still unclear. Previous studies confirmed a variety of miRNAs with abnormal expression in placental tissues are related to fetal growth and development (8-10). The results of Jiang et al. (11) showed miR-21 expression was up-regulated in NDFMS, which speculated miR-21 might be involved in the occurrence of NDFMS.

So far, there was no study on the patterns of miRNA expression in placental tissues of NDFMS. We screened out miRNAs with abnormal expression in NDFMS placental tissues through miRNA microarray, and studied the mechanism of key miRNA in this study.

## Results

### Clinical data

Clinical characteristics of the study population are summarized in Table 1. NDFMS and control were matched by maternal age, pre-pregnant body mass index, gestational weeks and infant gender distribution. The birth weight of neonates and weight gain during pregnancy were significantly higher in NDFMS than that in normal controls (*P* < 0.05).

**Table 1.** Clinical characteristics of the study population.

| Characteristic | Total (n = 91) | Control (n = 47) | NDFMS (n = 44) | *P*-value |
| --- | --- | --- | --- | --- |
| Maternal age (years) | 26.42 ± 2.87 | 26.15 ± 0.36 | 26.70 ± 0.48 | NS |
| Gestational age (weeks) | 39.71 ± 1.01 | 39.70± 0.15 | 39.72 ± 0.15 | NS |
| BMI before pregnancy (kg/m<sup>2</sup>) | 20.71 ± 3.32 | 20.05 ± 0.33 | 21.43 ± 0.62 | NS |
| Weight gain during pregnancy (kg) | 18.21 ± 4.77 | 16.60 ± 0.59 | 19.97 ± 0.75 | < **0.001** |
| Birth weight (g) | 3278.65 ± 516.49 | 3309.60 ± 47.01 | 4250.82 ± 41.41 | < **0.001** |
| Infant gender, n (%) |  |  |  | NS |
| Male | 43 (47) | 23 (49) | 20 (45) |  |
| Female | 48 (53) | 24 (51) | 24 (55) |  |

Values are mean ± SD. BMI, body mass index. NS = not significant.

### Results of miRNA microarray

We selected 18 cases of NDFMS and 18 subjects of control randomly, which mixed as one sample respectively in the miRNA microarray analysis. A total of 760 miRNAs were detected by miRNA microarray, and 264 and 318 miRNAs could be detected in placental tissues of normal control and NDFMS, respectively (Ct < 30). There were 21 miRNAs that expressed 4 times higher in NDFMS group, and 11 miRNAs expressed 8 times lower compared with controls.

### qRT-PCR validation in placentas

Twelve miRNAs (miR-331-3p, miR-204-5p, miR-106b-5p, miR-370-3p, miR-205-5p, miR-141-3p, miR-126-3p, miR-424-5p, miR-194-5p, miR-411-5p, miR-1291, and miR-1283) mentioned above were screened out from the perspective of bioinformatics, conservatism and practicality (Table S3) to examine the expression levels in placentas using qRT-PCR in others except those used in detection of miRNA microarray (29 of control, and 26 of NDFMS). The results showed the expression levels of miR-331-3p, miR-370-3p, miR-141-3p, miR-126-3p, miR-424-5p, and miR-411-5p were statistically different between NDFMS and normal controls, and *P* values were 0.047, 0.034, < 0.001, 0.023, 0.031, and 0.040, respectively (Fig. S2). Considering P-value and the fold change of expression, miR-141-3p was determined as the key miRNA (Fig. 1A), and fetal birth weight gradually increased along with the increased expression of miR-141-3p as the correlation analysis performed (P = 0.019, r^2^ = 0.101) (Fig. 1B), so we mainly explored the function of miR-141-3p in NDFMS.

**Figure 1.**
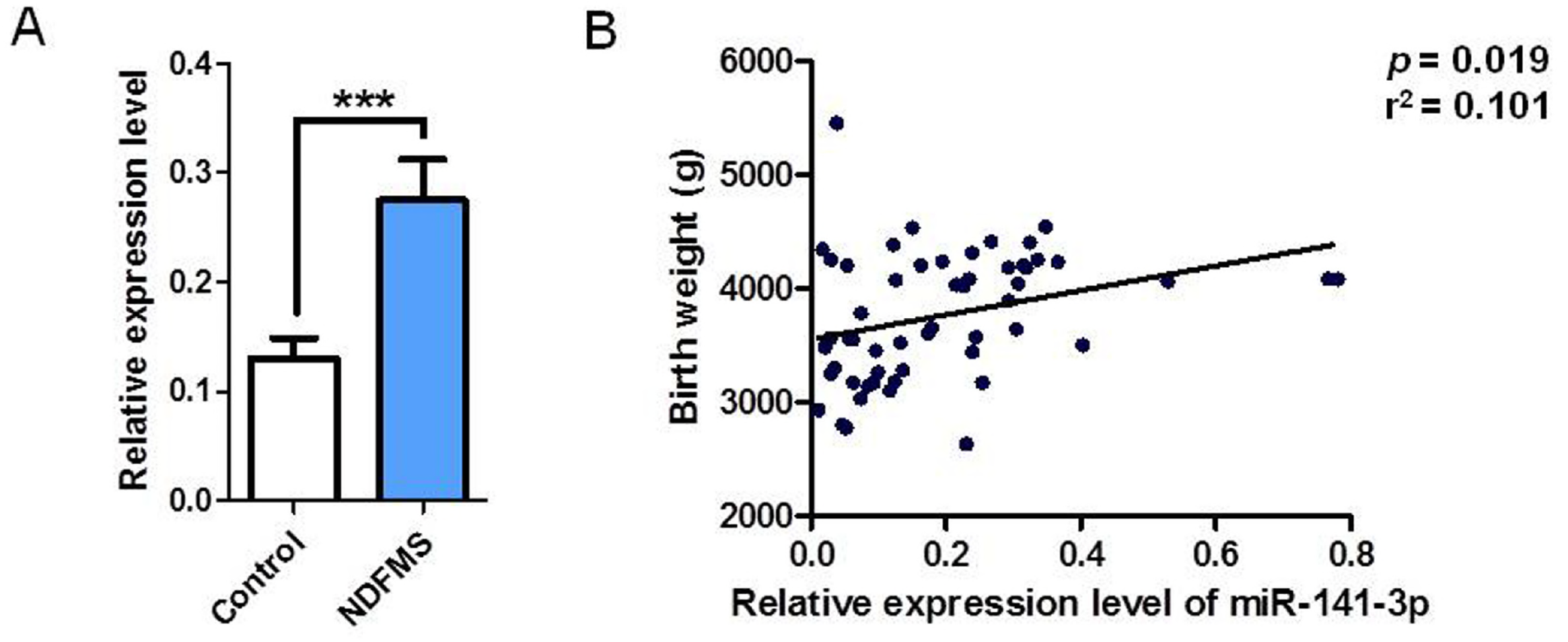
The expression level of miR-141-3p in placental tissues and the correlation with birthweight of fetus. **(A)** The expression level of miR-141-3p in NDFMS placental tissues compared with normal controls. **(B)** The correlation between expression level of miR-141-3p and fetal birth weight.

### Functional effects of miR-141-3p on proliferation, invasion, migration, apoptosis and cell cycle in HTR-8/SVneo cells

Cell Counting Kit-8 (CCK8) assays showed the proliferation was significantly increased in HTR-8/SVneo cells transfected with miR-141-3p mimics, whereas had no significant difference between inhibitors and controls at 48 h (Fig. 2A). Apoptosis assay and cell cycle assay showed there was no impact of miR-141-3p on cell apoptosis and cell cycle (Fig. 2B, 2C).

**Figure 2.**
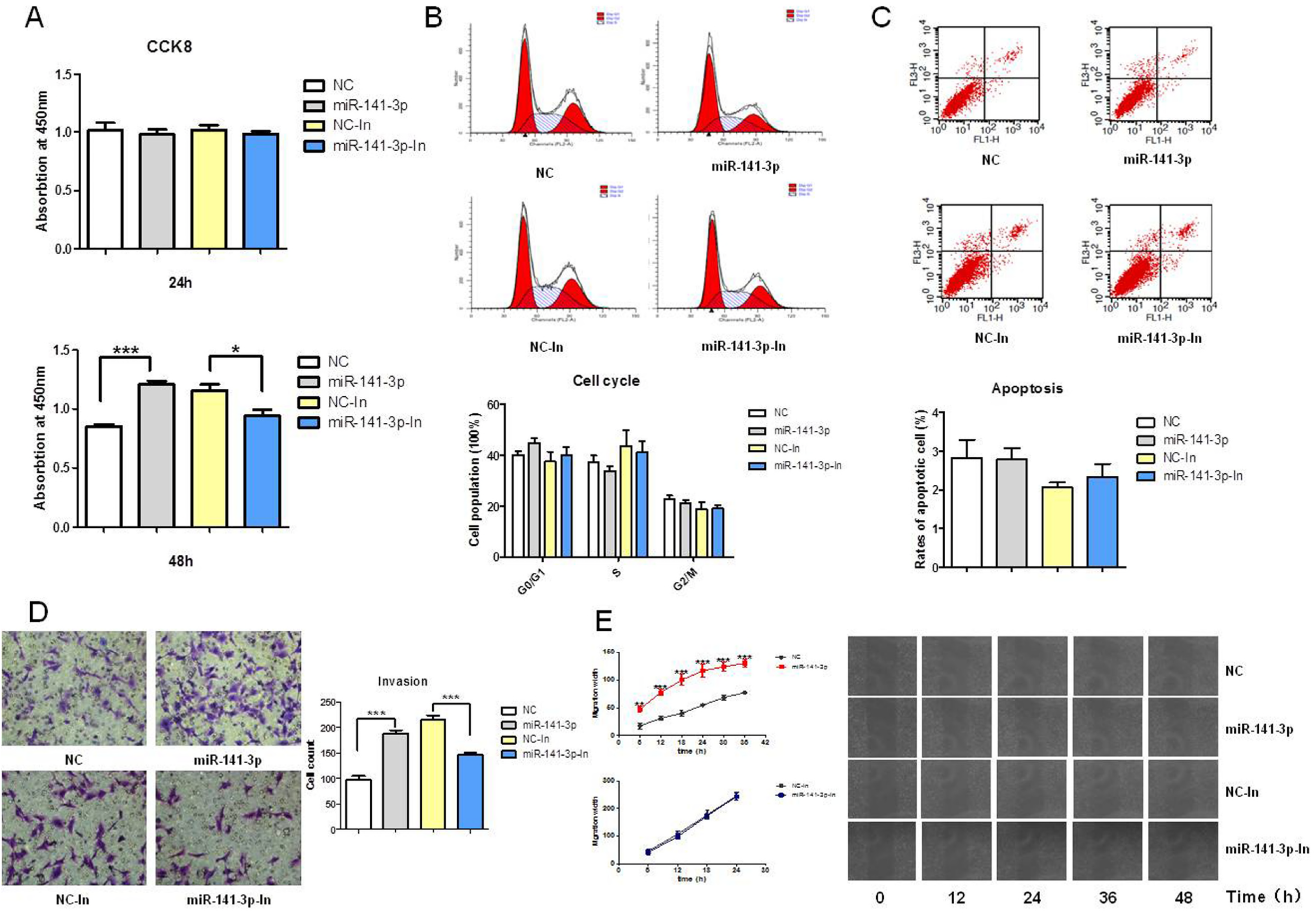
Effects of miR-141-3p on proliferation, cell cycle, apoptosis, invasion and migration of HTR-8/SVneo cells. **(A)** The proliferation ability of HTR-8/SVneo transfected with miR-141-3p mimic or inhibitor at 24 and 48 hours detected by CCK8. **(B)** Cell cycle of HTR-8/SVneo transfected with miR-141-3p mimic or inhibitor at 24 hours detected by flow cytometry. **(C)** Cell apoptosis of HTR-8/SVneo transfected with miR-141-3p mimic or inhibitor at 24 hours detected by flow cytometry. **(D)** The invasion ability of HTR-8/SVneo transfected with miR-141-3p mimic or inhibitor detected by crystal violet staining in transwell. **(E)** The migration ability of HTR-8/SVneo transfected with miR-141-3p mimic or inhibitor detected by BioSation IM-Q live cell workstation.

Transwell assays found miR-141-3p mimics increased the invasion of HTR-8/SVneo cells, while miR-141-3p inhibitors with the opposite result. Scratch assays showed miR-141-3p mimics enhanced migration ability while miR-141-3p inhibitors had no effect (Fig. 2D, 2E).

### Targeting *PLAG1* by miR-141-3p

In the prediction analysis, we identified 19 target genes of miR-141-3p. We examined the expression levels of the 19 genes (Fig. S3), and found only *PLAG1* was significantly decreased after transfected with miR-141-3p mimics (*P* < 0.001) (Fig. 3A). Western blot analysis showed miR-141-3p inhibited the expression of PLAG1 protein (Fig. 3B). Additionally, the expression of *PLAG1* gene in NDFMS placentas was significantly lower than that in controls (*P* = 0.007) (Fig. 4).

**Figure 3.**
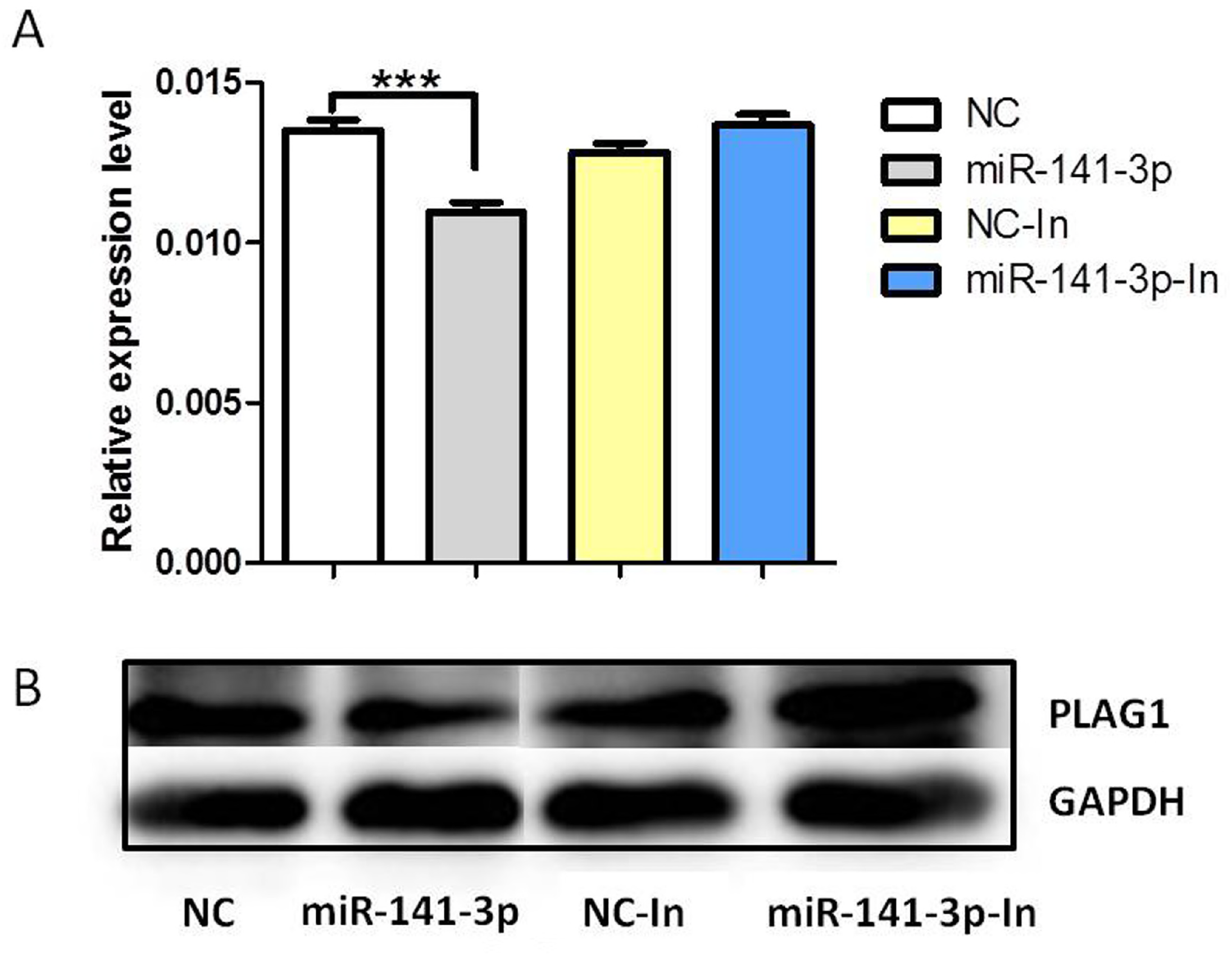
The expression level of PLAG1 mRNA and protein in HTR-8/SVneo cells after treated with miR-141-3p mimic and inhibitor. **(A)** Detection of *PLAG1* mRNA expression in miR-141-3p mimic or inhibitor-transfected HTR-8/SVneo cells by qRT-PCR, and normalized to GAPDH. **(B)** Detection of PLAG1 protein expression in miR-141-3p mimic or inhibitor-transfected HTR-8/SVneo cells by Western Blot, and normalized to GAPDH.

**Figure 4.**
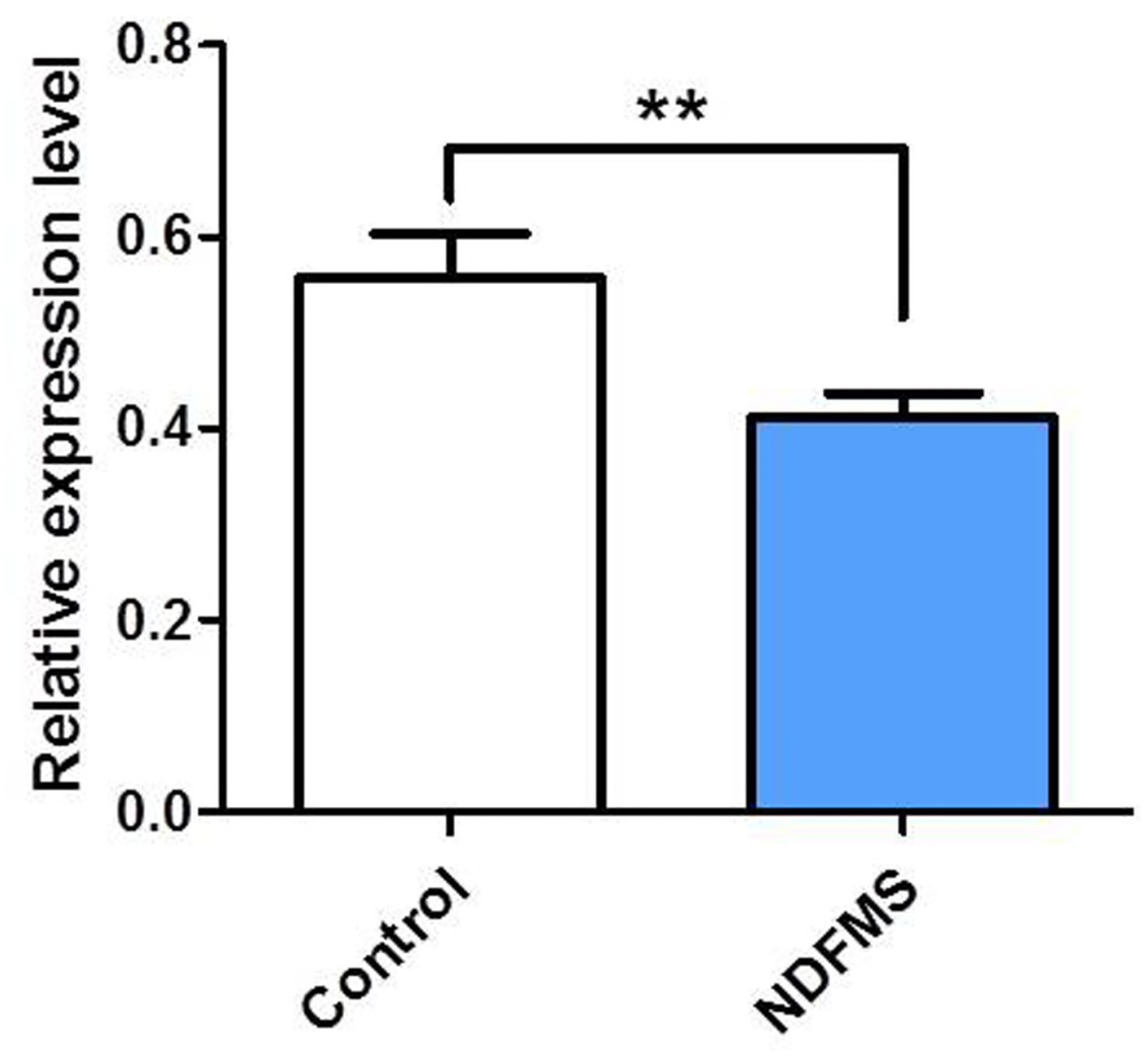
The expression level of *PLAG1* mRNA in NDFMS placental tissues compared with that of controls.

When overexpression of miR-141-3p and *PLAG1* at the same time, we found proliferation and invasion ability could be reversed by *PLAG1* on the basis of increased by miR-141-3p induced at 48 h (Fig. 5), which suggested miR-141-3p could involve in the regulation of trophoblast cell function by inhibiting the expression level of target gene *PLAG1*.

**Figure 5.**
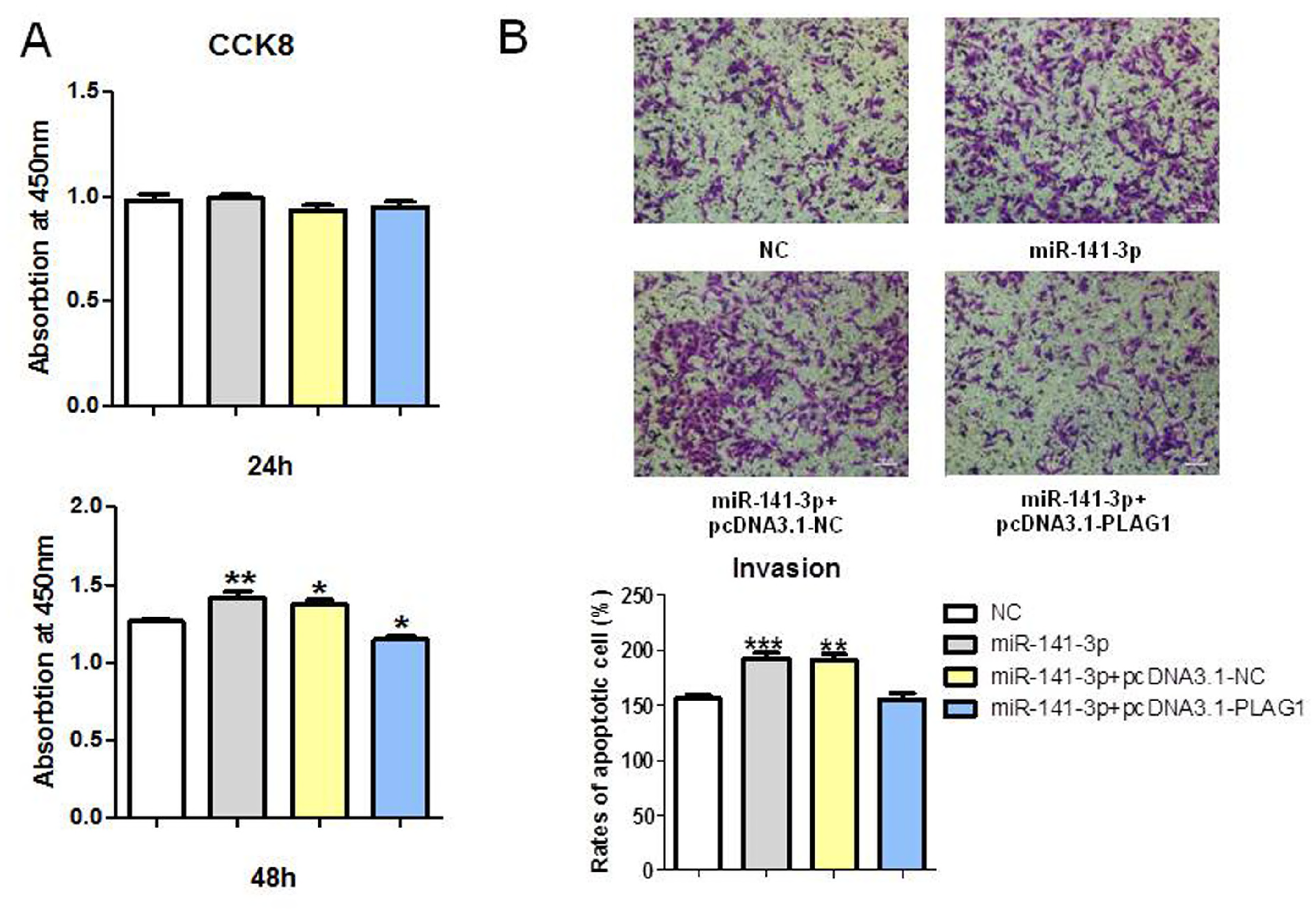
The proliferation and invasion of HTR-8/SVneo co-transfected with miR-141-3p and *PLAG1*. **(A)** The proliferation of HTR-8/SVneo co-transfected with miR-141-3p mimic and *PLAG1* overexpression plasmid at 24 and 48 hours. **(B)** The invasion of HTR-8/SVneo co-transfected with miR-141-3p mimic and *PLAG1* overexpression plasmid.

### Roles of miR-141-3p in birth weight of C57BL/6J mice

There was no difference between the birth weight of the treated group and controls in early pregnancy (Fig. 6), while the birth weight of the treated group was higher than controls in late pregnancy (*P* = 0.006) (Fig. 7). At the same time, there was no statistically significant difference about placental weight and diameter between treated group and controls neither in early pregnancy nor in late pregnancy (Fig. 6 and Fig. 7), as well as the body height of offspring (Fig. 6 and Fig. 7).

**Figure 6.**
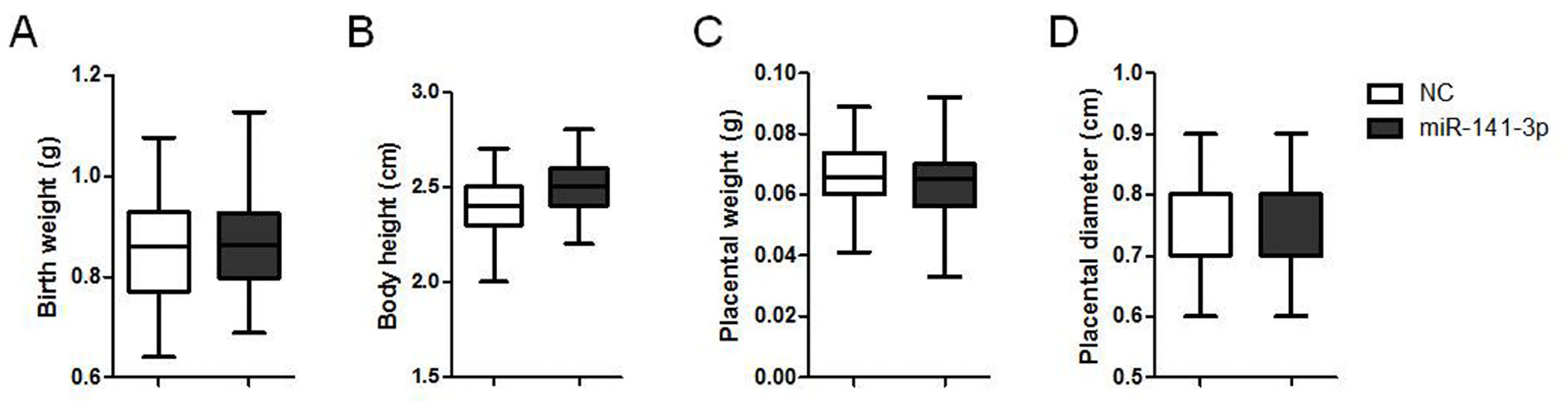
The effects of miR-141-3p treated at early pregnancy on offspring and placenta of pregnant C57BL/6J mice. **(A)** Birth weight of offspring. **(B)** Birth body height of offspring. **(C)** Placental weight. **(D)** Placental diameter.

**Figure 7.**
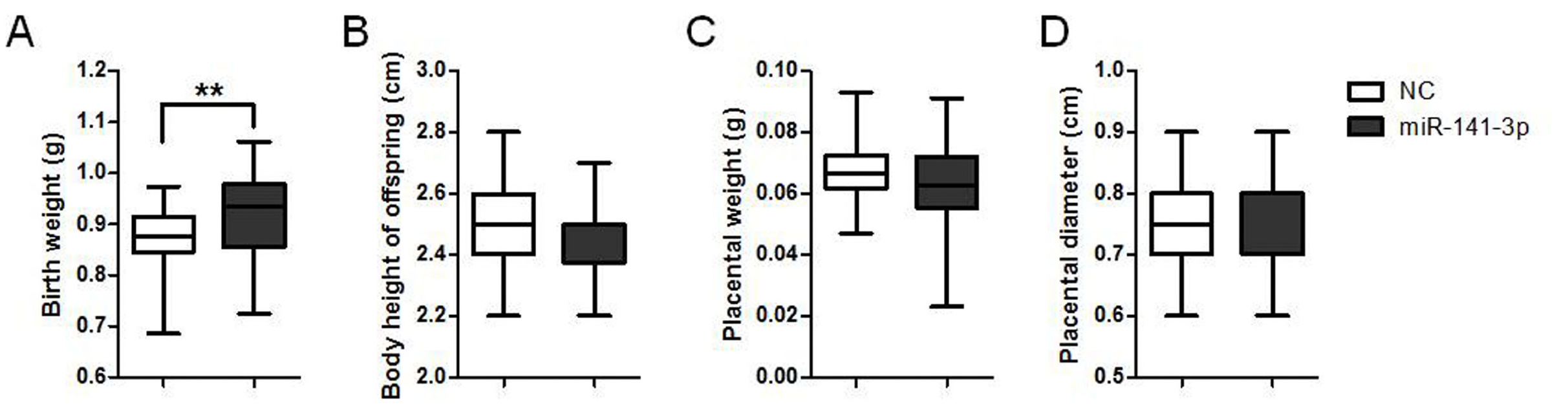
The effects of miR-141-3p treated at late pregnancy on offspring and placenta of pregnant C57BL/6J mice. **(A)** Birth weight of offspring. **(B)** Birth body height of offspring. **(C)** Placental weight. **(D)** Placental diameter.

## Discussion

We found the expression level of miR-141-3p was significantly increased in NDFMS placental tissues, suggesting aberrant expression of miR-141-3p may be related to the occurrence of NDFMS. miR-141-3p is a member of miR-200 family, which is highly conserved in vertebrates (12). The study of miR-200 family regulating fetal growth and development of population-based is rare, while relevant animal research is more common (13-15). Gerlinde et al. (15) found that expression levels of miR-200 family in the uterus and offspring (F3 generation) brain tissues changed when the parental generations (F0, F1 and F2) received stress stimulation at pregnant day 12 to 18. Lo et al. (16) detected known 157 miRNAs by miRNA microarray and found 17 miRNAs were 10 times higher in placental tissues than in maternal blood, which could not be detected in maternal blood after delivery. miR-141, miR-149, miR-299-5p and miR-135b were the four most abundant miRNAs in the placenta, in which the expression level of miR-141 in maternal blood significantly increased in the third trimester.

The cell experiments suggest miR-141-3p may participate in the development of NDFMS by affecting the proliferation and invasion of trophoblast cells. Ospina-Prieto et al. (17) confirmed the expression level of miR-141 in HTR-8/SVneo cells was significantly lower than that in normal and preeclampsia placenta. The expression level of miR-141 in normal placental tissues was 42394 times higher than that in HTR-8/SVneo cells. Markert et al. (18) also found the expression levels of miR-141 in all placental lineage cells were low, suggesting that the basal level of miR-141 in human chorionic trophoblast cells was very low, which may be the reason that the proliferation of HTR-8/SVneo cells treated by miR-141-3p inhibitor did not decrease as expected. Moreover, further studies are still needed to confirm this conclusion.

We found *PLAG1* is a target gene of miR-141-3p, which is a member of pleomorphic adenoma gene (PLAG) zinc finger protein family (19). Several studies found *PLAG1* is involved in the development of human cancers, such as salivary gland pleomorphic adenoma, adiponectomy, hepatoblastoma, and acute myeloid leukemia (20-22). In addition, a large number of studies have confirmed *PLAG1* plays an important role in the growth and development process, which can regulate the downstream target genes involved in cell proliferation process (23). We found *PLAG1* overexpression decreased cell proliferation, and could reverse the increase of cell proliferation and invasion induced by miR-141-3p, suggesting miR-141-3p regulates trophoblast cells function by inhibiting *PLAG1* expression, and thus participates in the development of NDFMS.

Because miR-141-3p affected proliferation and invasion ability of HTR-8/SVneo cells, we constructed the miR-141-3p overexpression model of placenta in early pregnancy and late pregnancy respectively. We purchased the reagent from Guangzhou RiboBio, which effect can be maintained for about three days, so as to ensure the treatment at early pregnancy will not be maintained until late pregnancy, which ensured the accuracy of modeling. We found fetal birth weight of treated group was higher only in late pregnant group, so we have reason to believe miR-141-3p may affect the proliferation at late pregnancy to participate in the occurrence of NDFMS, rather than influence the invasion at early pregnancy.

Previous studies reported the role of miR-141 in fetal growth (17, 24). Morales-Prieto et al. discovered miR-141 was highly expressed in placental tissues of preeclampsia, and overexpression of miR-141 enhanced the invasion ability of HTR-8/SVneo cells, which consistent with our findings. They also found miR-141 increased cell proliferation ability although the difference did not have statistical significance. And we found miR-141-3p was able to promote cell proliferation at 48 hours after treatment, possibly due to different experimental methods caused the different results. Markert et al. study also confirmed our results and they showed the reduction of miR-141 caused by leukemia inhibitory factors inhibited the proliferation of choriocarcinoma cell line JEG-3 (25). In addition, the study team also found miR-141 expression level in placental tissues at the late pregnancy was higher than the early pregnancy, may be due to placenta grows fastest along with the growth of fetus in late pregnancy, which also suggested miR-141 in placenta is related to proliferation function and confirmed the rationality of our results.

Our data suggested aberrantly elevated expression level of miR-141-3p may contribute to the pathogenesis of NDFMS by inhibiting the expression of downstream target gene *PLAG1*, thus affecting the proliferation in late pregnancy. Our study provided a foundation for explanation of miRNA changes related to NDFMS and expanded the current understanding of its pathogenesis. A summary of miR-141-3p mediated signaling model in NDFMS is shown in Fig. 8.

**Figure 8.**
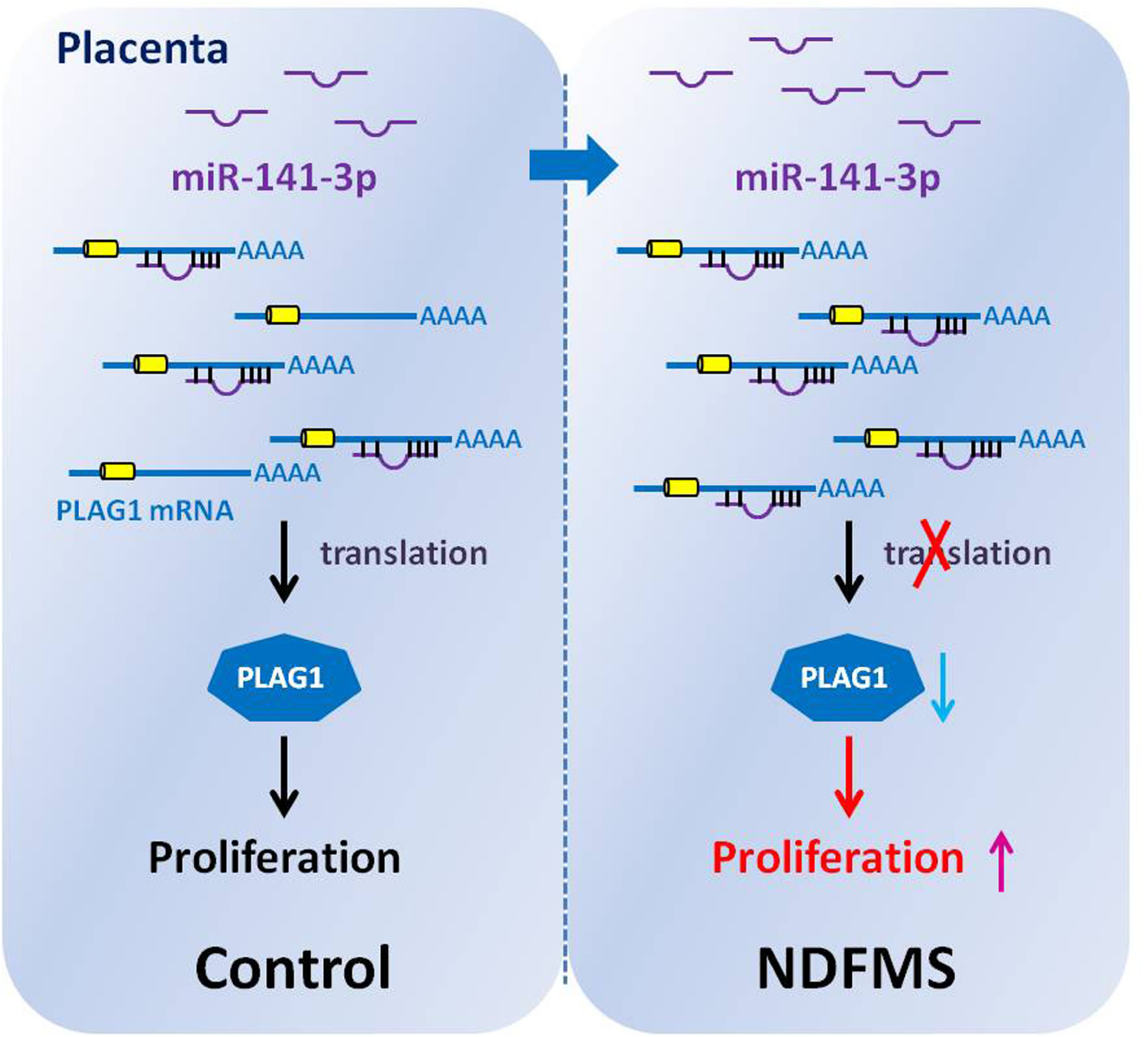
A model was presented for miR-141-3p mediated NDFMS. Up-regulation of miR-141-3p in placenta inhibits the translation of PLAG1, which drives macromosia by promote the proliferation of trophoblast cell that mediated the pathology of NDFMS.

## Materials and Methods

### Human material samples and ethics

A total of 91 subjects (47 normal controls, 44 non-diabetes macrosomia pregnant women) delivered in Changzhou Maternal and Child Health Hospital between September 2014 to June 2015 were included. Inclusion criteria of NDFMS: 1. Full-term birth (≥ 37 weeks and < 42 weeks), birth weight ≥ 4000 g; 2. Normal blood glucose and oral glucose tolerance test (OGTT) result, negative urine sugar; 3. Without various pregnant complications. At the same time, pregnant women with fetuses of full-term delivery and normal birth weight randomly were selected as a control group, and removed some with a variety of pregnancy complications.

The placental tissues were frozen in liquid nitrogen immediately after delivery, and then stored at −80°C. The study was approved by the Ethics Committee of Nanjing Medical University, and all pregnant women signed the informed consent before participation.

### RNA extraction, miRNA array analysis, and quantitative real-time PCR (qRT-PCR)

We used TRIzol Reagent (Invitrogen Life Technologies Co, CA, USA) to extract total RNA, and reverse transcribed using the specific Megaplex RT primers (Life Technologies). Thereafter, the expression level of 760 different miRNAs was performed using the TaqMan^®^ Array Human MicroRNA A+B Cards Set v3.0 (Life Technologies) on a 7900 Real Time PCR System (Applied Biosystems). Data were analyzed by DataAssist v3.0 (Life Technologies).

For qRT-PCR, PrimeScript^™^ RT reagent Kit (Takara Bio, Tokyo, Japan) was used to reverse transcribe RNA sample to cDNA, and PCR analysis was performed using SYBR^®^ Premix Ex Taq^TM^. GAPDH and RNU6 were analyzed as internal controls for mRNA and miRNA, respectively. The primer sequences for PCR are shown in Table S1 and Table S2.

### Cell culture and transfection

HTR-8/SVneo cell was cultured in RPMI 1640 medium containing 10% fetal bovine serum and 10% double antibody (100 U/ml penicillin and 100 U/ml streptomycin) at cell incubator of 37 °C and 5% CO_2_. We used the logarithmic growth phase cells to construct model by transfected miRNA mimic or inhibitor (GenePharma, Shanghai, China) using transfection reagent Lipofectamine 2000 (Invitrogen Corp, CA, USA). Then, we used these cells to conduct experiments after 24 h or 48 h according to the experiment needs.

### Cell proliferation assays

CCK-8 (Dojindo, Kumamoto, Japan) was used to measure cell proliferation ability. Then measured the absorbance at a wavelength of 450 nm by TECAN infinite M200 Multimode microplate reader (Tecan, Mechelen, Belgium).

### Cell cycle analysis

We collected the treated cells, washed with cold PBS, fixed in cold 70% ethanol overnight at −20 °C. PI (Sigma, MO, USA) was used to stain the fixed cells for 30 min at room temperature under dark condition. Cell cycle distribution of stained cells was analyzed with FACS Calibur Flow Cytometer afterwards, and fraction of the cell cycle was measured by Modfit LT version 3.0 software.

### Apoptosis assays

We harvested the treated cells, washed twice with PBS, and then processed cells using Annexin V-FITC Kit (BD Biopharmingen, NJ, USA). The cells were analyzed using FACS Calibur Flow Cytometer immediately after incubated with Annexin V-FITC.

### Cell invasion analysis

The treated cells were transferred to the transwell chamber with 100 μl empty RPMI 1640 medium, while the 24-well plate contained 600 μl complete medium as the container of the transwell chamber (26). After 24 h, the transwell chamber was transferred to 95% methanol to fix cells for 30 min, then stained by crystal violet for 30 min, washed thrice with PBS. After drying, we captured the pictures by light microscope, and used the cell number in vision as a measure.

### Cell migration analysis

We used 10 μl pipette tip to draw a straight line in each hole of 6-well plate with treated cells, and washed twice with PBS. Next, the BioSation IM-Q live cell workstation was used to record the migration situation of cells to the middle real-timely and capture dynamic change graphs. Then we used Image-Pro Plus software to analyze the pictures, which take the horizontal distance of cells migrated as the quantitative indicator.

### miRNA targets prediction

In order to find out the potential target genes of miR-141-3p, we collected the target genes from TargetScan (http://www.targetscan.org/vert_72/), miRDB (http://mirdb.org), RNA22 (https://cm.jefferson.edu/rna22/Interactive/), PicTar (https://pictar.mdc-berlin.de/cgi-bin/PicTar_vertebrate.cgi), PITA (https://genie.weizmann.ac.il/pubs/mir07/mir07_prediction.html) and chose the genes at least in four software as the target genes.

### C57BL/6J mice feeding and experiment

We used miR-141-3p agomir (micro*ON*^TM^ miR-141-3p agomir) (RiboBio, Guangzhou, China) to construct miR-141-3p overexpression model in placental tissues via tail vein, and used micro*ON*^TM^ agomir negative control treated controls. Eight-week-old pregnant C57BL/6J mice were divided into two groups: early pregnant group (treated: control = 5: 5) and late pregnant group (treated: control = 7 : 5). Early pregnant group mice were treated twice at pregnant day 5 and day 8, while late pregnant group mice were treated twice at pregnant day 14 and 17. Then, they were sacrificed at pregnant day 18, and measured fetal weight. Besides, we treated some unpregnant mice twice with 3 days interval (number of treated: control = 4 : 5), and sacrificed at 10 days after the second treatment, and observed weight fluctuation (The results of effect on body weight and food intake were shown in Fig. S1).

### Statistical analysis

Data were analyzed by SPSS 18.0 and presented with GraphPad Prism 5.01. If data were in normal distribution, the student’s t-test was used to compare two groups. Otherwise, the Mann-Whitney U-test were used. All results are expressed as mean ± standard error (X ± S.E.) without special instructions. *P* < 0.05 was considered statistically significant. *P* < 0.05: *, *P* < 0.01: **, *P* < 0.001: ***.

## Acknowledgements

Supported by National Natural Science Foundation of China (814012138), Jiangsu Provincial Medical Youth Talent (QNRC2016110), “333 Project” Science Research Project of Jiangsu Province (BRA2016197), “six big talent peak” Project of Jiangsu Province (WSN-283), Clinical Medicine Center Project of Nantong City (HS2016005), Jiangsu Overseas Visiting Scholar Program for University Prominent Young & Middle-aged Teachers and Presidents, and the Priority Academic Program for the Development of Jiangsu Higher Education Institutions (Public Health and Preventive Medicine).

## Author contributions

W.W., Q.T., L.C., D.G., and H.J. conceived and designed the experiment. D.G., H.J., Y.C., J.Y., Z.F., J.L., and X.H. performed the experiments. D.G., X.W., and W.W. performed the bioinformatics analysis. D.G., W.W., Q.T., L.C., Y.X., and X.W. wrote and revised the manuscript.

